# Combinatorial bioassay for fast screening of organic agrivoltaic materials

**DOI:** 10.64898/2026.05.15.725514

**Authors:** Margalida Vidal-Tur, Martín Martín-Ruiz, Alfonsina Abat Amelenan Torimtubun, Mariano Campoy-Quiles, Jaume F. Martínez-García

## Abstract

Agriphotovoltaics (APV) combines crop production with solar energy generation to address increasing demands for food and energy while reducing land-use competition. Unlike conventional opaque photovoltaic systems, semitransparent organic photovoltaics (OPVs) selectively absorb light, potentially improving efficiency but also altering both light quantity and spectral quality, key factors affecting plant growth. Here, we developed a rapid bioassay based on hypocotyl elongation to evaluate plant responses to OPV-filtered light using *Arabidopsis thaliana* and *Cardamine hirsuta*, two species with contrasting shade strategies. Screening a diverse set of OPV materials revealed that plant growth responses depend more on spectral composition than on total light intensity alone. Certain materials, such as PTB7-Th and D18, produced growth patterns similar to neutral shading, while others promoted elongation. Our analyses identified blue light wavelengths, linked to cryptochrome activity, as more critical than red light wavelengths, linked to phytochrome activity, for maintaining normal development. These findings provide a scalable framework to assess OPV-plant compatibility and demonstrate that optimizing spectral quality alongside light intensity is essential for designing efficient APV systems that sustain crop performance while generating renewable energy.

## INTRODUCTION

Agriphotovoltaics, agrovoltaics or agrivoltaics (APV) refers to the coexistence of land use for agriculture and solar photovoltaic electricity production. Sharing the same piece of land for both activities can reduce or/and eliminate the competition of agricultural land in the current context of an increasing demand of both food and energy production. Current commercial silicon-based photovoltaic solar panels have a broad absorption range, extending from the ultraviolet (UV) to the near-infrared (near-IR, nearly 1200 nm), which makes them fully opaque. Therefore, in the design of conventional APV systems, the elevation, the spacing between rows and tilt angles of opaque solar panels are adjusted to reduce shade and allow solar radiation to reach the ground under the solar PV structure for crop growth (Abidin et al., 2021; Ghosh, 2023). While several large-scale studies have shown increased land use for opaque-based APV (Macknick et al., 2022), it has been pointed out that the use of semitransparent photovoltaics can enhance further the land use due to a reduce spectral competition (Emmott et al., 2015).

Plant photosynthesis is a light-dependent process that depends on their photosynthetic pigments (chlorophylls and carotenoids), that absorb primarily in the so-called photosynthetic active radiation (PAR) region (400-700 nm). This coincides in part, although not completely, with the visible light spectrum for humans. The amount of PAR light is normally expressed by the Photosynthetic Photon Flux Density (PPFD, µmolꞏm^-2^ꞏs^-1^). On the other hand, the sun delivers light from a much broader spectrum (from ca 300 nm to 2500 nm after atmospheric filtering) than the PAR. Therefore, the use of semi-transparent photovoltaic materials in which the photovoltaic device absorbs light in a complementary fashion to plants has emerged as a promising strategy to circumvent the limitations associated with opaque modules, and, moreover, help to simplify the design of the APV structures (Emmott et al., 2015). These materials should allow the light transmitted beneath the electricity-producing solar cells (i.e., the portion of the spectrum not used for electricity generation) to be sufficient to sustain photosynthesis and exceed the light compensation point, which is the light intensity where the rate of photosynthesis (that results in CO_2_ incorporation) exactly matches the rate of cellular respiration (that results in CO_2_ loss) (Valladares & Niienemets, 2008). If these conditions are matched, transmitted light will permit crop thriving.

From the different semiconductors that can be used to make solar cells, organics are, arguably, the best suited for the task due to the specific mechanisms behind their absorption profile. Specifically, the molecular nature of the organic semiconductors typically results in a strong absorption band above the bandgap, associated to the main singlet transition and the vibronic sidebands, followed by a valley of minimal absorption before rising again at higher energies (often blue/UV) due to absorption bands associated to electronic transitions of more localized electrons. Besides their specific absorption profile, organic photovoltaics (OPVs) are highly efficient, approaching 21% (Chen et al., 2025; Zhang et al., 2025) (cf. dye sensitized PV). They can be produced on flexible substrates, and do not contain highly toxic materials (such as lead, Pb).

In contrast to APV systems using opaque PV that reduce the intensity of the light for all wavelengths, the transmission of light using semi-transparent OPV materials is not neutral, i.e., they absorb specific spectral regions, potentially altering light quality. This is very relevant, as plants use light for obtaining energy through photosynthesis, but also as a source of environmental information. As sessile organisms, plants need to respond and acclimate to the changes in their environment, including changes in the light quantity (amount) and quality (light spectrum). To do so, plants have evolved a battery of light receptors (photoreceptors) that detect changes in the quantity and quality of the light and trigger optimal development changes for plant survival or/and thriving in natural settings. In particular, changes in light quality inform about the close presence of competing vegetation (Casal, 2012; Martinez-Garcia & Rodriguez-Concepcion, 2023).

In natural and agronomic environments, plants may be found either quite isolated (low vegetation density) or, more commonly, growing close together (high vegetation density). High density can limit light availability and compromise plant development. To cope with these light gradients, plants have evolved two main alternative strategies: shade avoidance and shade tolerance. Shade avoidance involves a series of growth and development adjustments aimed at “escaping” shaded conditions (such as promotion of hypocotyl and/or stem elongation) even if it comes at the cost of reduced photosynthesis or defense capabilities. In contrast, shade tolerance is more common in plants adapted to low-light or heavily shaded environments. These plants adopt a conservative growth strategy to survive under limited light (Valladares & Niienemets, 2008; Martinez-Garcia & Rodriguez-Concepcion, 2023). Importantly, these two strategies illustrate the diverse way plants have evolved to thrive under varying light quantities.

*Arabidopsis thaliana*, a dicot herbaceous plant from the *Brassicaceae* family, has been developed as the main model system for shade-avoidance research (Martinez-Garcia & Rodriguez-Concepcion, 2023). This small, self-pollinating weed with a short life cycle and generous seed production, offers extensive genetic tools, resources and techniques to explore the genetic and molecular mechanisms of many plant responses. The findings obtained with such plant can usually be extended to plant species of commercial interest which might have more complex growth requirements and are generally harder to manipulate in the lab. On the other hand, *Cardamine hirsuta*, a close relative to *A. thaliana* with a contrasting shade-tolerant growth habit, has also been developed as a versatile experimental system, as it shares many of the advantages for lab use that could be found in *A. thaliana*. As a shade tolerant plant, *C. hirsuta* displays a conservative growth and resource use strategy, and low phenotypic plasticity to alter growth, such as reduced elongation in response to vegetation proximity, that is reduced or absent compared to shade-avoiding species (Molina-Contreras et al., 2019; Paulisic et al., 2019).

OPV installations may reduce the amount of available light by partially absorbing solar radiation (light quantity) and alter the light spectrum (light quality). This creates complex light environments that can cause unexpected developmental or architectural changes in crops, potentially decreasing productivity (i.e., amount of crop produced per input, including land, water, nutrients, labor, or per growing season) and yield (i.e., total amount of crop produced per unit area). Therefore, it is key to understand how exposure to different light qualities resulting from OPV-filtered light environments impacts on plant development, photosynthesis and/or architecture, parameters that ultimately might impact plant production.

While several proof-of-concept studies have shown promising results, a general method to evaluate compatibility of materials and plant growth would help to take educated decisions. In this work, we aim to systematically analyze the compatibility of OPVs with plant growth by advancing a combinatorial-like bioassay for the fast evaluation of the impact of a range of OPV materials in a small-size format on seedling development. As plant systems, we selected *A. thaliana* and *C. hirsuta*, two small-size and related species with divergent strategies to vegetation proximity. In terms of OPV systems, we chose active materials with largely varying bandgaps, as well as absorption full width at half maximum for the main absorption band. We analyzed the growth of the two plants under OPV filters of different active layer thickness and collected a large dataset that allowed us to evaluate statistical correlations between plant growth, biological spectra and OPV features.

## RESULTS

### SECTION 1: PRESENTATION OF THE METHOD

#### A general method to evaluate compatibility of OPV materials and plant growth is missing

One of the hypotheses when designing novel OPV materials for APV is that they should minimally impact the photosynthetic relevant light (i.e., the PAR) to reach plants. Different figures of merit have been used for this. From the photovoltaic side, the average visible transmittance (AVT) has been often used. Recently, the average light transmission in the PAR region, *aka* average photosynthetic transmittance (APT) has also been proposed (Stallknecht et al., 2023). Both of these figures of merit assume that all wavelengths within the PAR are equivalent. However, this is not the case and efficiency exhibits two peaks in the rate of photosynthesis occurring in the blue (400-500 nm) and red (600-700 nm) parts of the spectrum, with the green (500-600 nm) contributing less to plant growth. The action spectrum of the plant is the magnitude that expresses such efficiency (McCree, 1972). Other figures of merit, such as the overlap between the action spectrum, the transmission of the OPV and the illumination, have also been proposed (Emmot et al., 2015). In addition, there are several reported average action spectra, as well as other biologically relevant spectra that have been used to design OPVs for AGV, including the absorption profiles of the two chlorophylls (McCree, 1972).

To contextualize the diversity of experimental approaches and biological metrics used to evaluate plant responses under spectrally modified light environments, we compiled a summary of representative studies in the field (**Table 1**). These studies go from controlled laboratory experiments to greenhouse and field-based agrivoltaics system, highlighting the wide range of plant species, experiment duration, light sources and filters, and evaluation criteria employed. Plant responses to spectrally modified environments have been evaluated using a wide range of biological metrics, including morphological traits (e.g., stem or hypocotyl elongation, leaf area), biomass accumulation, physiological parameters as photosynthetic activity or chlorophyll content, and fruit yield (**Table 1**). However, the diversity of experimental conditions, plant species and evaluation criteria makes it difficult to directly compare results across studies.

**Table 1.**
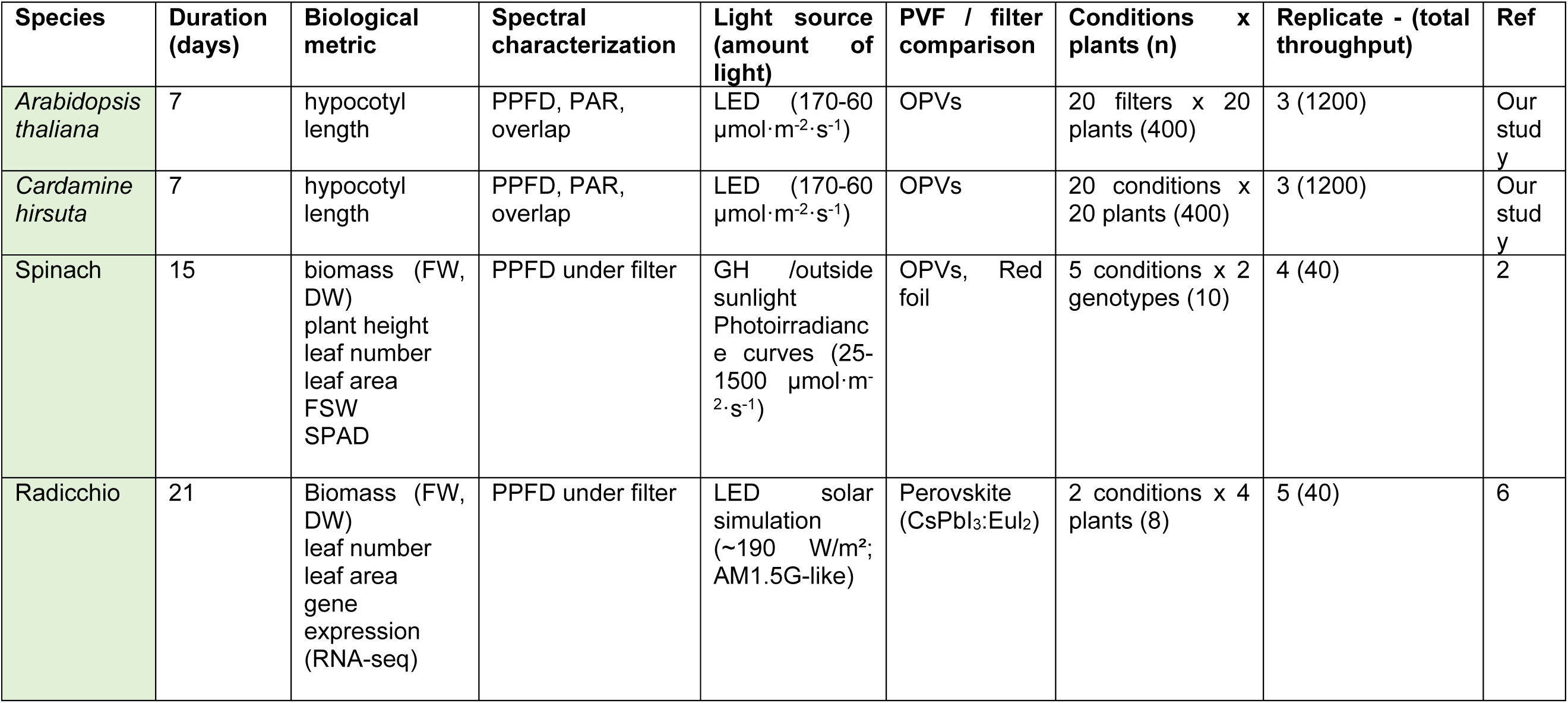
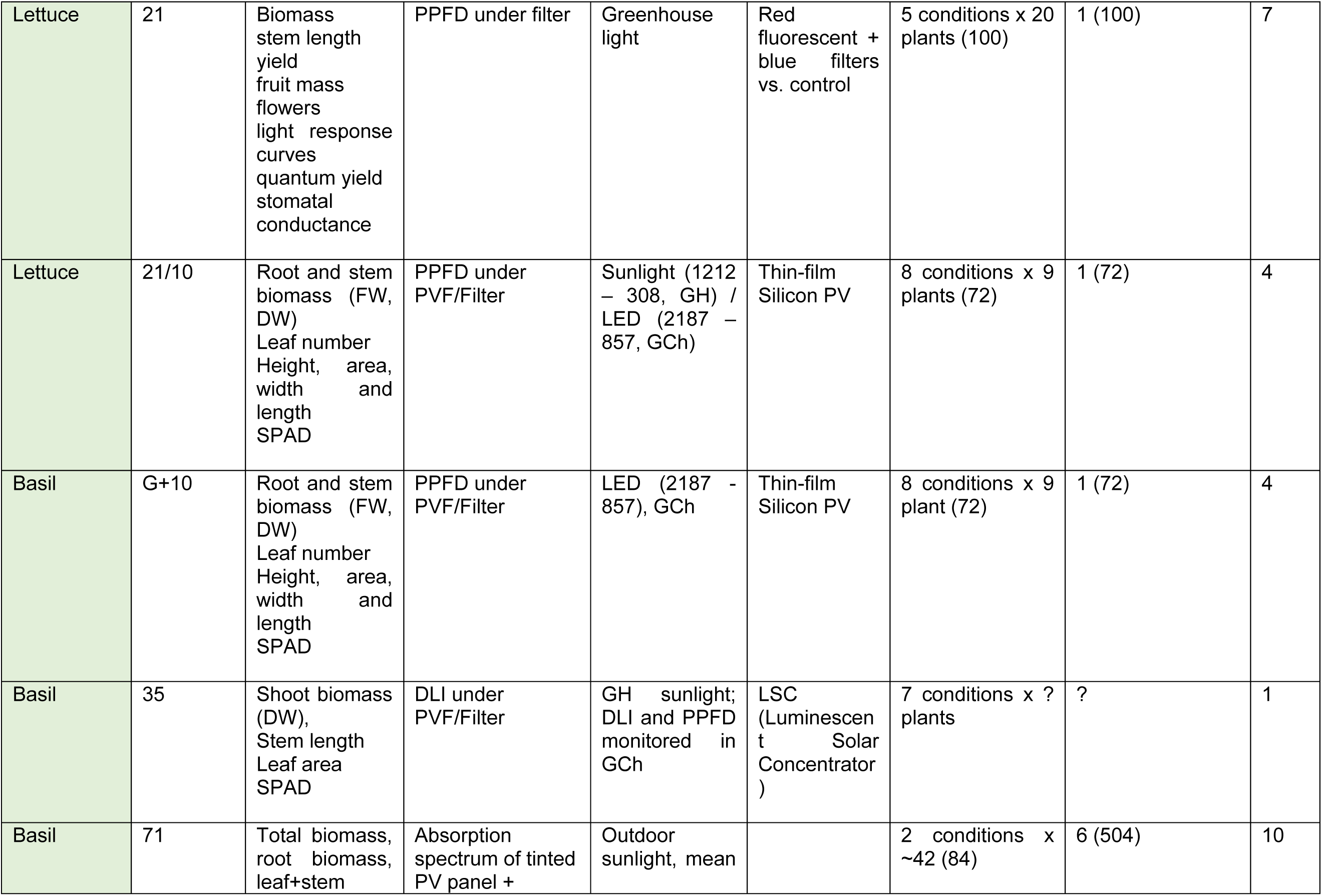
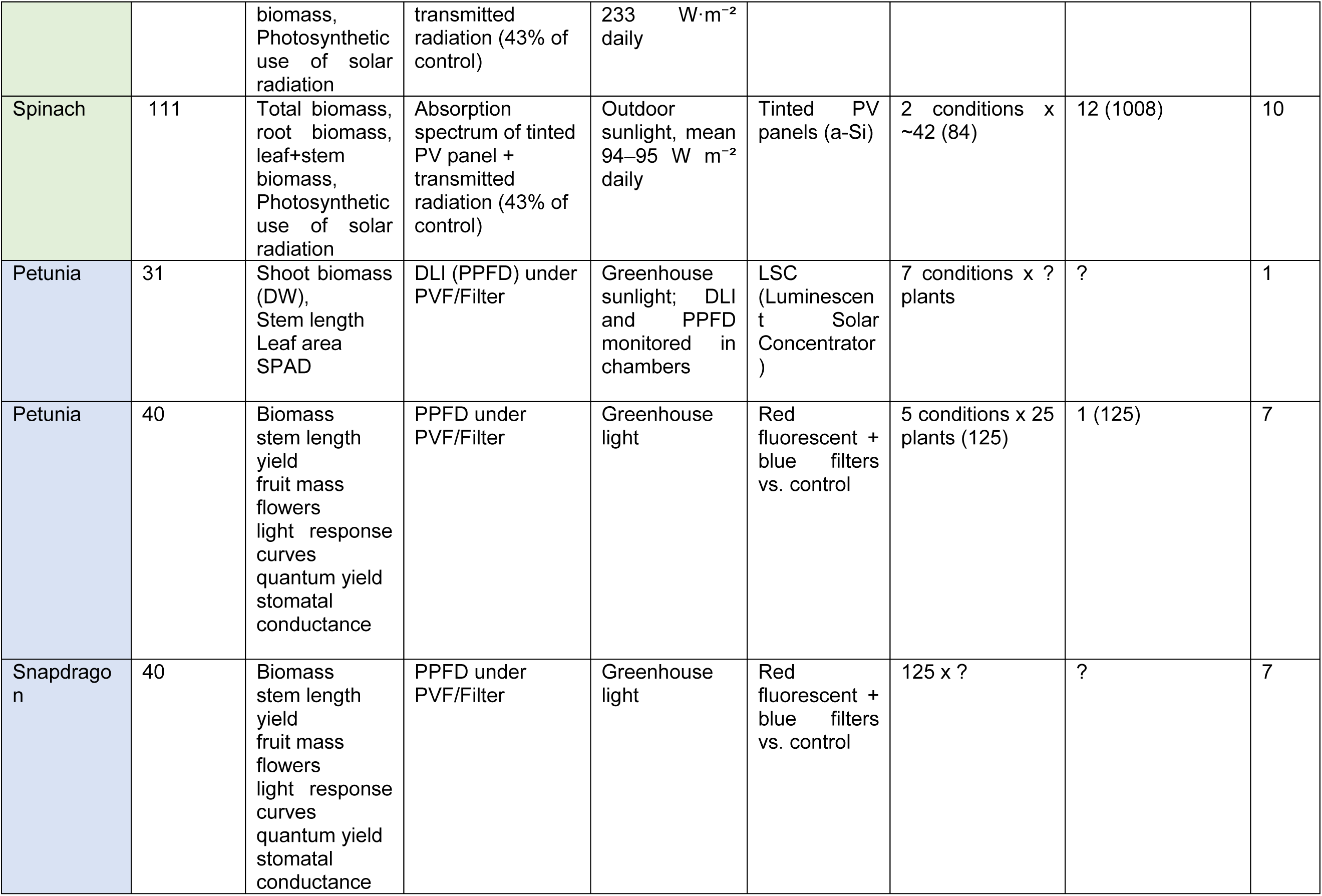
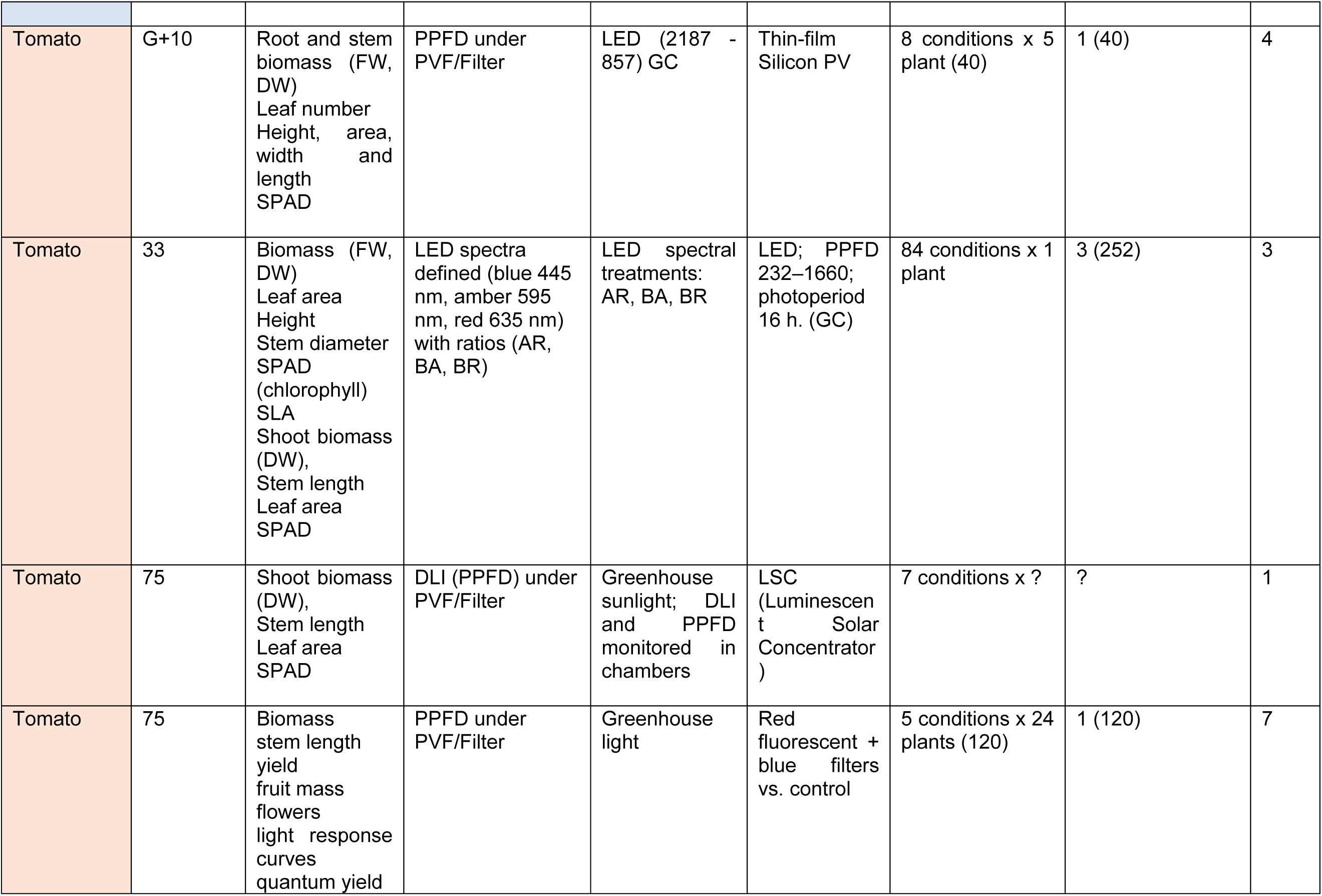
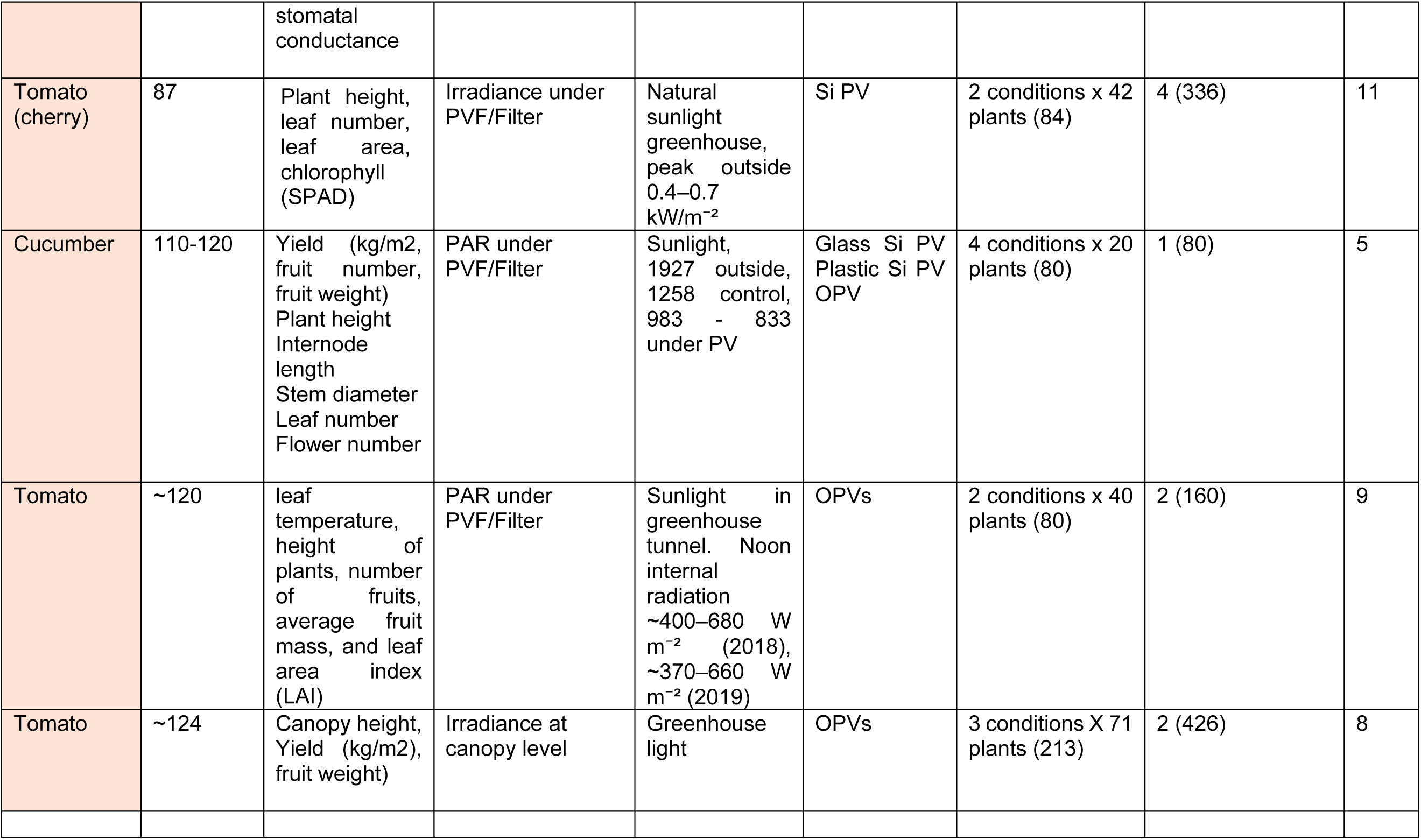
Summary of experimental studies evaluating plant responses under spectrally selective photovoltaic or optical filtering systems. The table includes representative examples from controlled laboratory experiments, greenhouse studies, land field-based agrivoltaic systems. For each study, key parameters are reported: plant species, experimental duration, biological metrics used to assess plant performance (e.g., morphology, biomass, physiological responses), level of spectral characterization, light source and intensity conditions, type of photovoltaic (PVF) or optical filtering system, and number of treatments. Abbreviations: GCh (growth chamber); GH (greenhouse); G, germination time; DLI (Daily Light Integral); FW, fresh weight; DW, dry weight; FSW, fresh leaf weight; SPAD, Soil Plant Analysis Development (SPAD) chlorophyll meter; SLA, Specific leaf area, which is the ratio of leaf area to leaf dry mass.

Morphological traits, such as stem or hypocotyl elongation and leaf area, are among the most used metrics to evaluate plant response under spectrally modified light environments. These parameters are relevant as they are highly sensitive to light quantity and quality. In contrast, the characterization of the light environments in these studies is typically based on integrated metrics such as PPFD or DLI (Daily Light Integral) measured under the different PV/filter system. While these parameters provide a measure of the total light availability, they assume that all wavelengths within the PAR region contribute equally to plant performance. However, this has not been tested before.

#### Hypocotyl length of *A. thaliana* and *C. hirsuta* is a rapid and effective parameter to monitor seedling growth under a range of light conditions

First, we wanted to set a bioassay to measure and quantify the impact of a series of OPV materials on plant growth. In order to make a fast evaluation of a large number of OPV materials and thicknesses, we chose to explore the growth of seedlings of *A. thaliana* and *C. hirsuta*, whose growth is affected by light quantity and quality (Martinez-Garcia & Rodriguez-Concepcion, 2023). Seeds of these two species were germinated and grown under increasing amounts of white light (W) for a week. As a control, seedlings were also grown in the dark for the same time period. Then, we monitored three morphological traits known to respond to light: hypocotyl length, cotyledon area and distance between cotyledons, as a proxy of cotyledon longitudinal expansion (**Figure 1**). While hypocotyl elongation decreased as the light that seedlings were growing under was increased, cotyledon area and distance between cotyledons increased. More importantly, the differences in hypocotyl elongation between the treatments were always greater than the differences between cotyledon parameters in both species, indicating that this response displayed the highest dynamic range (ratio between the maximum and minimum response). Therefore, in the following set of experiments the bioassay was established by measuring hypocotyl length.

**Figure 1.**
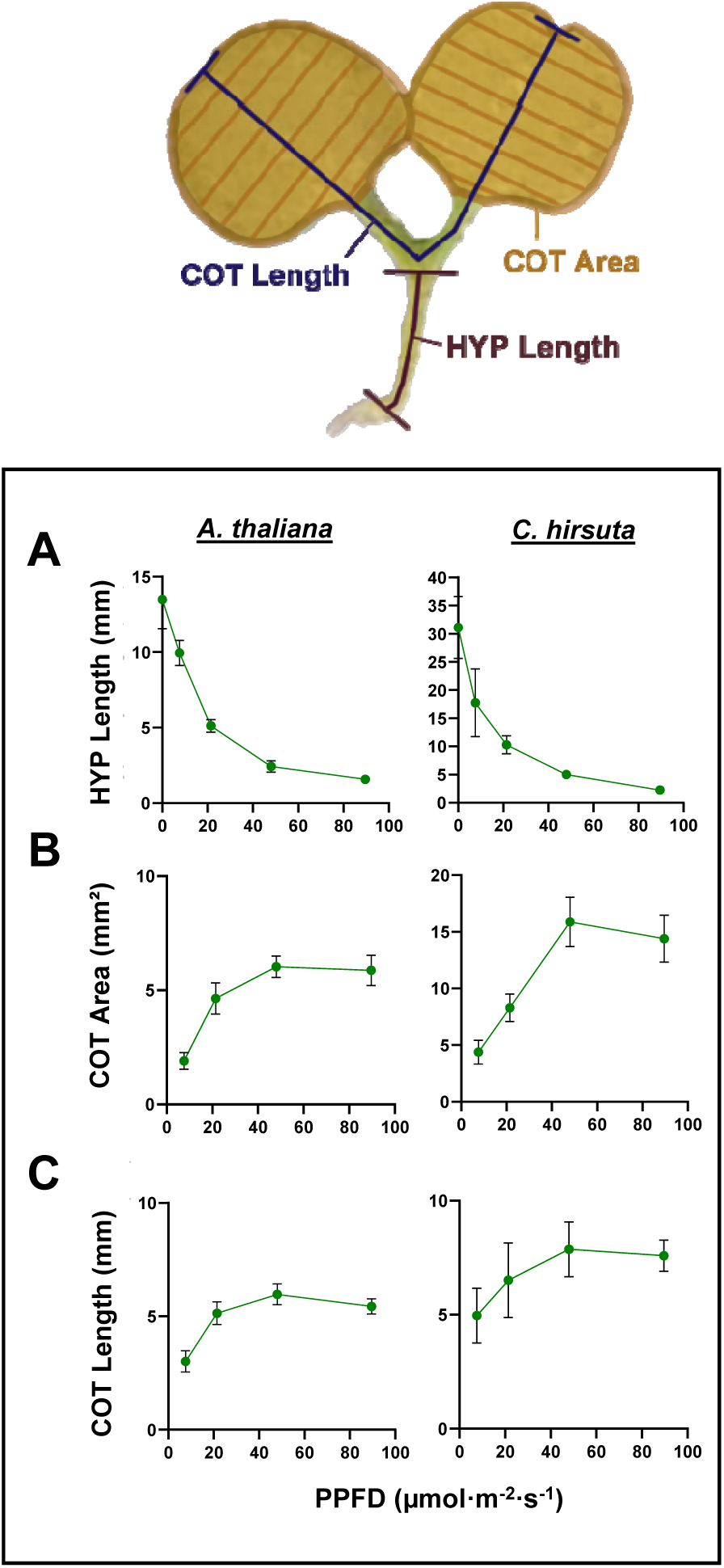
Morphological phenotype of *Arabidopsis thaliana* and *Cardamine hirsuta* seedlings when germinated under progressively lower amounts of light. **(A)** Scheme of the different parameters of a seedling scored as morphological responses to light conditions: hypocotyl length, as the distance of this organ, cotyledon area, as , and . Seedlings of *A. thaliana* and *C. hirsuta* were grown for 7 days under different light intensities of W (R:FR > 2.3). The different light intensities were obtained using increasing layers of filtering paper, that work as a neutral filter, to reduce the amount of light they received. Graphs represent **(A)** hypocotyl length, **(B)** sum of the area of the two cotyledons of each seedling and **(C)** distance between cotyledons through the main axis relative to the Photosynthetic Photon Flux Density (PPFD). Values of hypocotyl length are the mean ± SD of 30 seedlings. Values of cotyledon area are the sum of the mean cotyledon area of both cotyledons of at least 10 seedlings. Cotyledon length corresponds to the distance… Values of cotyledon length are the mean ± SD of at least 10 seedlings. Values of PPFD are the mean ±SD of three independent measurements taken at the beginning of the experiment at the exact placement of each box.

#### Hypocotyl assays suggest that *C. hirsuta* tolerate lower light intensities than *A. thaliana*

Next, hypocotyl length was measured in *A. thaliana* and *C. hirsuta* seedlings grown for 7 days under increasing amounts of light of high (white light, W) and low red (R) to far-red light (FR) ratio, R:FR (W enriched in FR, W+FR) (**Figure 2**). Changes in the R:FR are detected by the phytochrome photoreceptors, which have a central role in detecting the proximity of vegetation that might compete for resources and eventually result in light starvation (Casal, 2012; Martinez-Garcia & Rodriguez-Concepcion, 2023). As before (**Figure 1**), hypocotyls of *A. thaliana* and *C. hirsuta* seedlings were shorter the higher the quantity of light they had grown under. Additionally, *A. thaliana* seedlings grown under W+FR displayed longer hypocotyls than those W-grown, a behavior not observed in *C. hirsuta* seedlings, whose hypocotyls responded similarly under both, W and W+FR (Martinez-Garcia & Rodriguez-Concepcion, 2023). This is consistent with the divergent shade habit of these two species. However, under low amounts of light (PPFD < 15-20 µmolꞏm^-2^ꞏs^-1^), *A. thaliana* hypocotyls stopped responding differentially to W and W+FR. More importantly, these comparisons indicated that the *C. hirsuta* elongated relatively less in response to the same amount of light than *A. thaliana* (**Figure S1**), in agreement with its shade-tolerant growth habit (Martinez-Garcia & Rodriguez-Concepcion, 2023).

**Figure 2.**
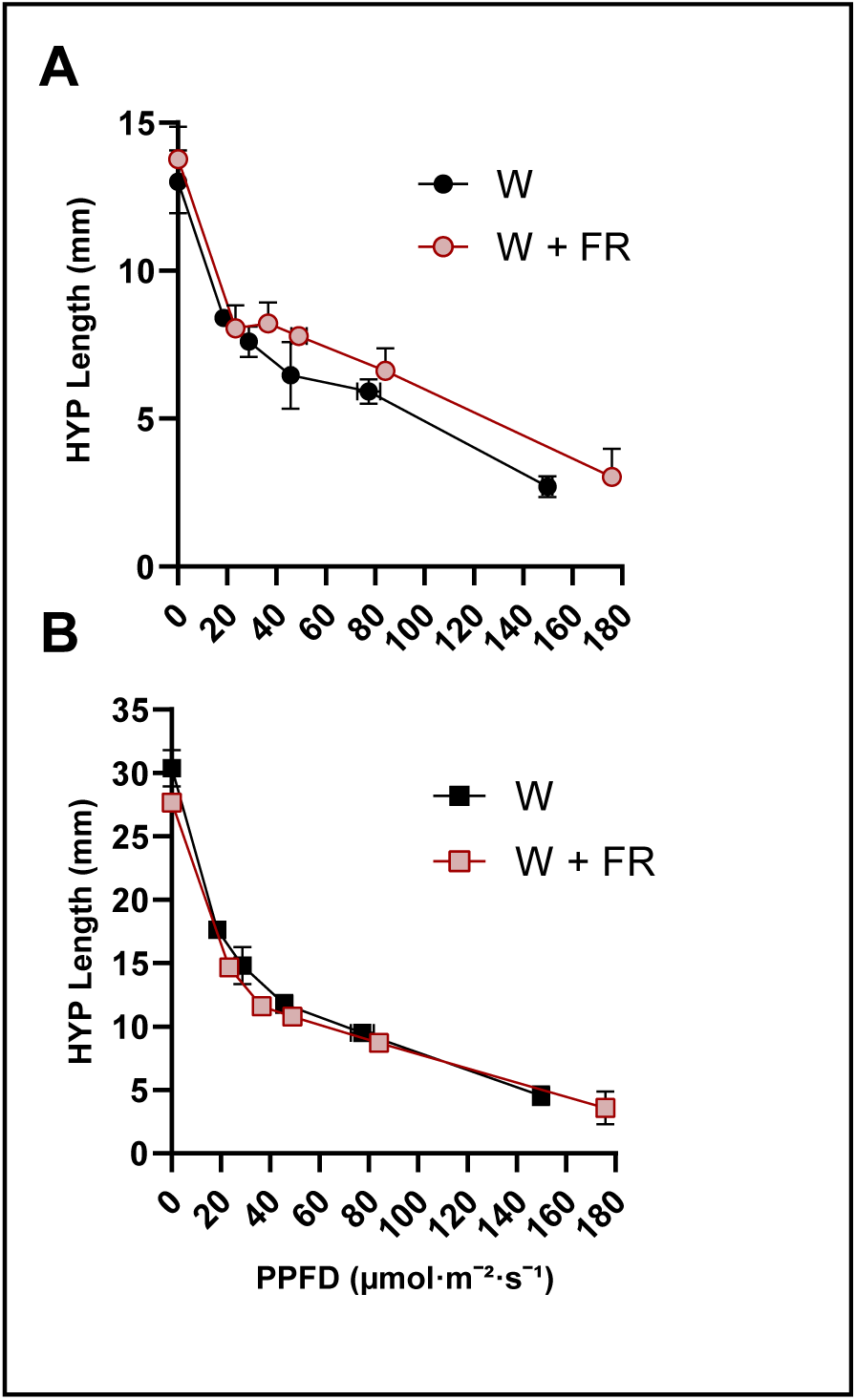
Differential hypocotyl elongation response of *Arabidopsis thaliana* and *Cardamine hirsuta* to different light quantity and quality conditions. Seedlings of. **(A)** *A. thaliana* and **(B)** *C. hirsuta* were grown for 7 days under different light intensities of W (R:FR > 2.3) or W+FR (R:FR ∼ 0.05). The different light intensities were obtained using increasing layers of filtering paper, that work as a neutral filter, to reduce the amount of light they received without altering the light spectrum. Graphs represent hypocotyl length (y axis) relative to the Photosynthetic Photon Flux Density (PPFD) (x axis). Values of hypocotyl length are the mean ±SD of three independent biological replicates. Each biological replicate is the mean of ≈ 18 seedlings. Values of PPFD are the mean ±SD of three independent measurements taken for each replicate at the beginning of each experiment.

### SECTION 2: EVALUATION OF DIFFERENT OPV MATERIALS

After evaluating the approach with neutral filters on two plant species, we next use the developed method to measure the effect of different OPV materials on the growth of the selected plants. Specifically, we started with the donor material, which typically overlaps more strongly with the PAR, then we evaluated the acceptors and, finally, selected blends. This stepwise approach enabled the identification of material combinations compatible with plant growth while generating a comparative dataset for later correlation analyses.

To carry out these studies, we fabricated a series of PET-based filters using different organic semiconductor materials varying the thickness of this layer, and thus the overall transmission. This produced a library of semitransparent filters with a range of spectral and optical characteristics (**Figure 3A**).

**Figure 3.**
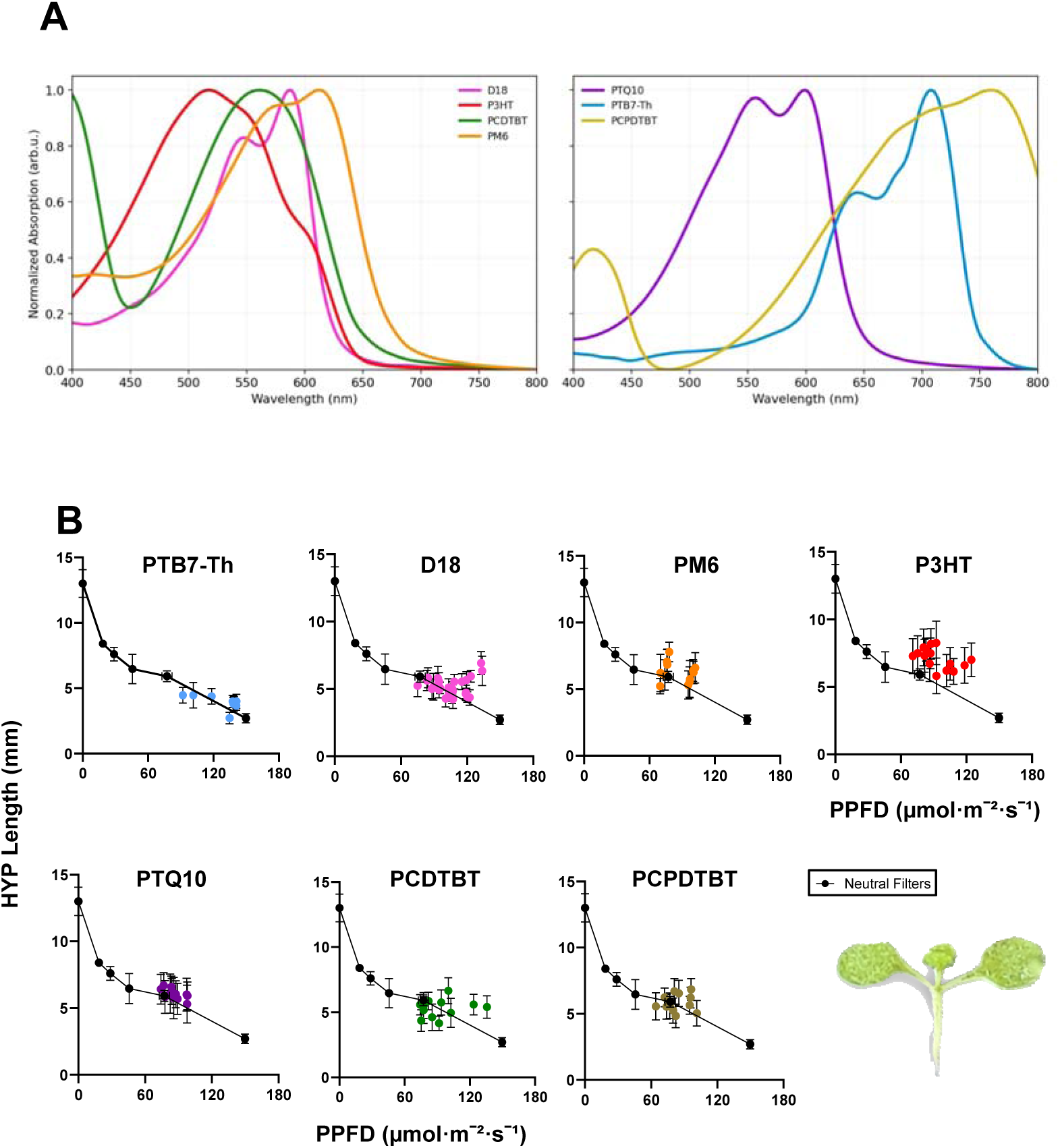
Hypocotyl elongation response of *A. thaliana* seedlings under different donor filters. Seedlings of *A. thaliana* and were grown for 7 days under filters made of different photoactive donor materials **(A)** showing different absorption spectra. **(B)** At the end of the experiments hypocotyl length was represented relative to the Photosynthetic Photon Flux Density (PPFD) transmitted by the different donor filters analyzed: PTB7-Th, D18, PM6, P3HT, PTQ10, PCDTBT and PCPDTBT. Graphs represent Values of hypocotyl length are the mean ±SD of three independent biological replicates. Each biological replicate is the mean of 12-20 seedlings. Values of PPFD correspond with three independent measurements taken for each replicate at the beginning of each experiment. Results are plotted over the results obtained from seedlings of the same species grown under the same conditions but using neutral filters to reduce the amount of light they grew under without changing the light spectrum (“Neutral Filters”), which are used a reference response under non-selective (neutral) filters.

#### Light filtered by PTB7-Th and D18 materials appear to affect less plant growth than neutral filters

We evaluated the impact of the light filtered by seven different donor materials by growing seedlings of *A. thaliana* and *C. hirsuta* for 7 days (**Figures 3, 4**). For each material, we fabricated between 3-5 filters with different film thickness. The absorption spectra of the analyzed donor materials (**Figure 3A**) illustrate that each material absorbs light preferentially in specific wavelength regions, generating distinct spectral filtering effects. Consequently, the light transmitted through each filter differs not only in total intensity but also in spectral composition. The resulting hypocotyl length values were evaluated against the PPFD resulting from different donor materials and thicknesses, and compared to the data for neutral filters, that reduce the quantity of light without affecting light spectra. These comparisons indicated that, out of the seven tested materials, PTB7-Th and D18 were the ones under which hypocotyls elongated less (i.e., the most similarly to those growing under neutral filters for a similar or even smaller PPFD). Seedlings grown under PCDTBT and PCPDTBT filters showed a greater dispersion of their hypocotyl lengths relative to the neutral filters. Seedlings grown under PTQ10, PM6 and P3HT filters showed longer hypocotyls than under neutral filters (**Figure 3**). The same tendency could be observed when growing *C. hirsuta*, although differences in hypocotyl length between seedlings grown under neutral filters and under the different OPV material filters were less pronounced. Interestingly, in this case PTB7-Th was more efficient in inhibiting hypocotyl elongation than D18 (**Figure 4**).

**Figure 4.**
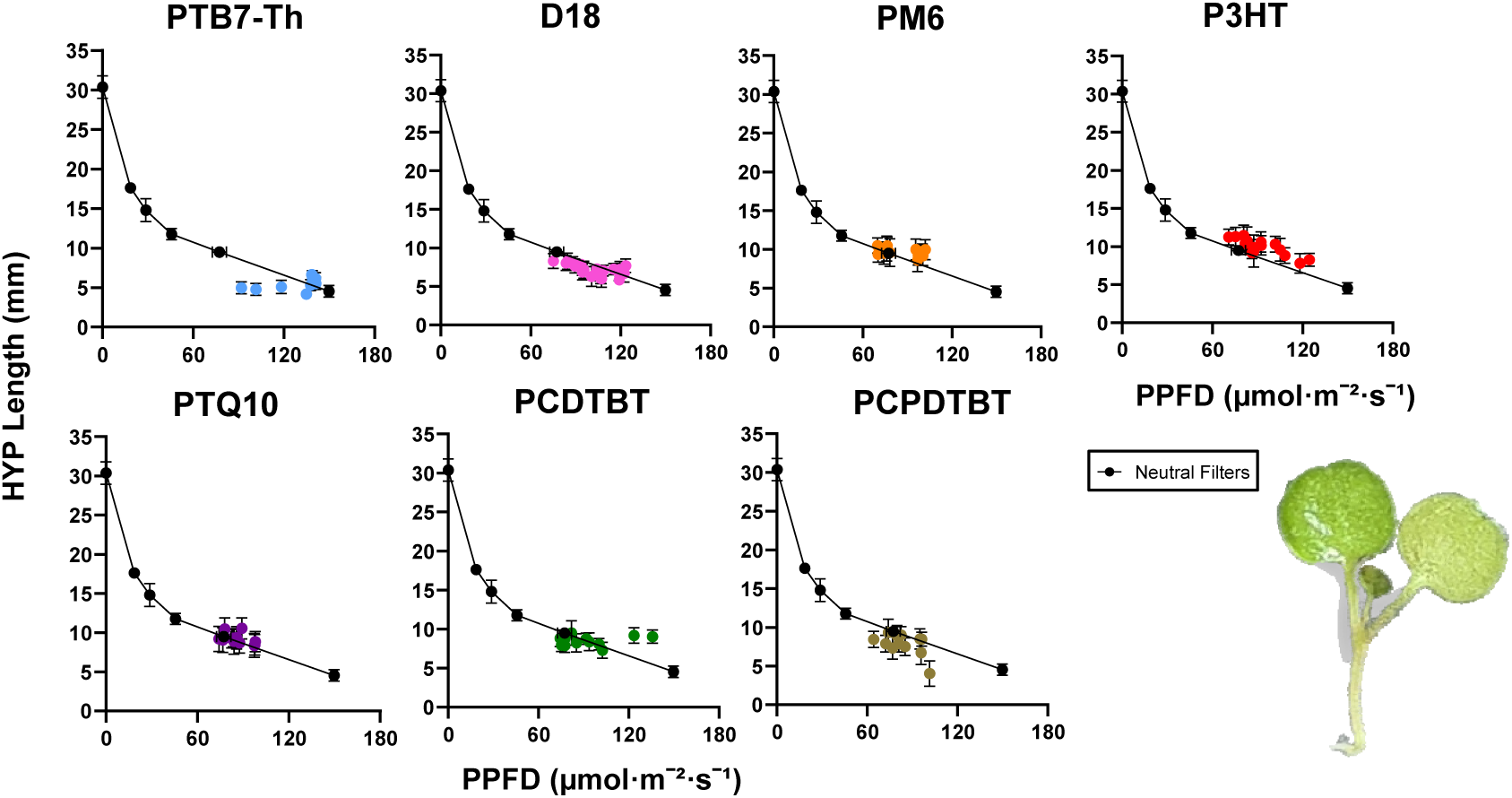
Hypocotyl elongation response of *C. hirsuta* seedlings under different donor fiters. Seedlings of *C. hirsuta* were grown for 7 days under filters made of different photoactive donor materials showing different absorption spectra (as seen in Figure 1A). At the end of the experiments hypocotyl length was represented relative to the Photosynthetic Photon Flux Density (PPFD) transmitted by the different donor filters analyzed: PTB7-Th, D18, PM6, P3HT, PTQ10, PCDTBT and PCPDTBT. Graphs represent Values of hypocotyl length are the mean ±SD of three independent biological replicates. Each biological replicate is the mean of 12-20 seedlings. Values of PPFD correspond with three independent measurements taken for each replicate at the beginning of each experiment. Results are plotted over the results obtained from seedlings of the same species grown under the same conditions but using neutral filters to reduce the amount of light they grew under without changing the light spectrum (“Neutral Filters”), which are used a reference response under non-selective (neutral) filters.

These results indicate that changes in spectral quality, and not only transmitted PPFD, play an important role in seedling growth responses and thus AVT or APT might not be the most suitable descriptors for agrovoltaics (see also below). Donor materials such as PTB7-Th, which shows stronger absorption in the green and red regions, and D18, with the peak of absorption in the green, were more effective at limiting hypocotyl elongation than materials that absorb mainly in the blue region, despite transmitting similar amounts of PPFD. These trends suggest that specific wavelength ranges differ in their biological relevance for growth regulation. As a result of the above analysis, PTB7-Th was selected as the best donor material for subsequent blend optimization.

#### Selection of the acceptor – impact of the donor to acceptor ratio on the Power Conversion Efficiency (PCE)

Having identified PTB7-Th as the most suitable donor material, we next focused on selecting an acceptor for blend fabrication. Based on the donor screening result and considering the need to preserve transmittance of as much PAR as possible for plant growth, a nice strategy was to select acceptor materials with absorption shifted toward longer wavelengths, outside the PAR region. In contrast to donor materials, acceptors offer greater potential to extend OPV absorption into the red (500-600 nm) and near-infrared region (> 700 nm). However, acceptor selection also requires consideration of electronic compatibility with the donor as well as the best PCE. i.e., the percentage of input irradiation that is converted into output power. In a recent study, we looked at the photovoltaic performance of PTB7-Th blended with 9 different acceptors, namely, Y6, IEICO-4F, BTP-eC9, Coi8DFIC, O-IDTBR, PC70BM, COTIC-4F, ITIC-4F, SiOTIC-4F, which resulted in PCEs of 10.45%, 10.32%, 9.38%, 9.06%, 8.84%, 8.48%, 8.38%, 8.12%, and 7.42%, respectively (Torimtubun et.al, 2024). Comparative optical and device analysis indicated that IEICO-4F provided a favorable balance between spectral and photovoltaic compatibility, since it gave almost the same PCE as Y6 and a comparatively red shifted absorption. For this reason, IEICO-4F was selected as the acceptor material for subsequent blend optimization studies.

#### Hypocotyl length is relatively insensitive to donor to acceptor ratio

Having selected PTB7-Th and IEICO-4F as the donor and acceptor materials, we next investigated whether adjusting the donor:acceptor ratio could improve the biological compatibility while preserving the photovoltaic potential of the blend. Previous work showed that reducing the proportion of PTB7-Th increased the device transparency, since PTB7-Th absorbs more in the PAR region than IEICO-4F (Martin-Trillo et al., 2026).

We tested the impact of the ratio of PTB7-Th:IEICO-4F on hypocotyl length of *A. thaliana* and *C. hirsuta* seedlings under two different light regimes: W (high R:FR), which simulates sunlight, and W+FR (low R:FR), which simulates the proximity of nearby vegetation (**Figures 5, 6**). We used five different ratios in which the amount of IEICO-4F (acceptor) was increased and PTB7-Th (donor) was reduced. Under the lower ratio blends (0.6:1.5 and 0.8:1.5) that were more transparent (transmitted higher values of PPFD), hypocotyl length of W-grown *A. thaliana* seedlings was slightly longer than the neutral filters. Interestingly, under the higher ratios of donor:acceptor blends (0.9:1.5 to 1.5:1.5) that were less transparent (transmitted reduced values of PPFD, so less light amount reached the seedlings) hypocotyl length was similar to that of seedlings under neutral filters. Under W+FR, the filtered light provided by the various blends resulted in hypocotyl lengths even shorter than those of seedlings grown under similar PPFD provided by neutral filters. This was particularly obvious in the 0.6:1.5 and 0.8:1.5 blends (**Figure 5**), indicating that these ratios attenuated the shade-inducible impact of low R:FR environments. The same tendency could be observed when growing *C. hirsuta*, although differences in hypocotyl length between seedlings grown under neutral filters and all the different blends were more advantageous (**Figure 6**). This suggests that combinations of OPV materials that minimize distortion of wavelengths in the blue range (i.e., those controlled by the cryptochrome photoreceptors) appear more compatible with maintaining normal seedling development.

**Figure 5.**
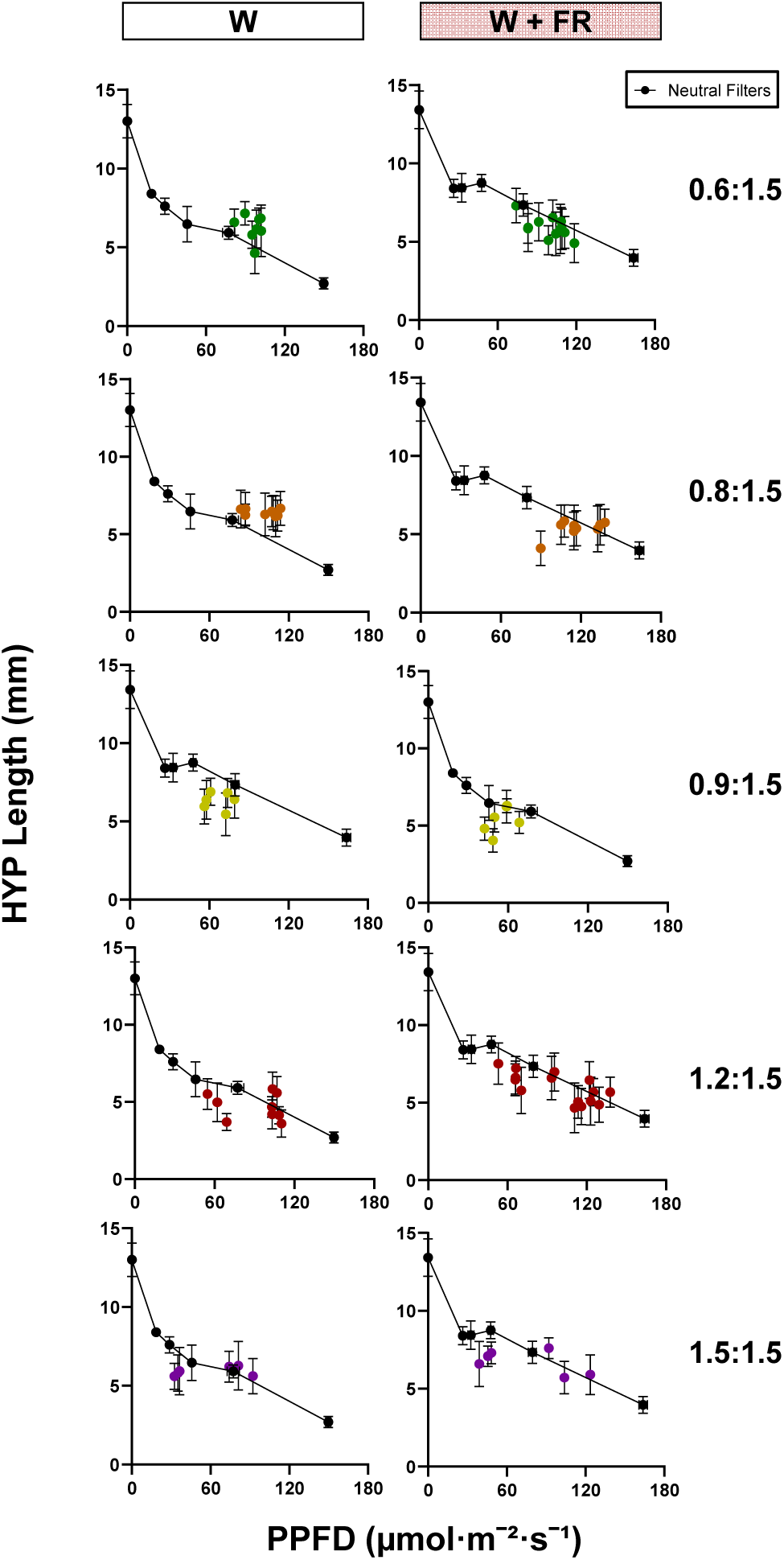
Hypocotyl elongation response of *Arabidopsis thaliana* under PTB7 – Th x IEICO 4F blended filters with different ratios of donor to acceptor material. – Hypocotyl elongation does not increase when increasing donor to acceptor ratio. Seedlings of *Arabidopsis thaliana* were grown for 7 days under filters made by blending PTB7 – Th (donor) x IEICO 4F (acceptor) materials in different ratios of donor to acceptor: **(a)** 0.6:1.5, **(b)** 0.8:1.5, **(c)** 0.9:1.5, **(d)** 1.2:1.5 and **(e)** 1.5:1.5. Graphs represent hypocotyl length (y axis) relative to the Photosynthetic Photon Flux Density (PPFD) (x axis). Values of hypocotyl length are the mean ±SD of three independent biological replicates. Each biological replicate is the mean of at least 15 seedlings. Values of PPFD correspond with three independent measurements taken for each replicate at the beginning of each experiment. “No Filter” control dots represent the results obtained from plants grown under the same experimental conditions but without a filter. “Blank” control dots represent the results obtained from plants grown under the same experimental conditions but using a transparent blank filter made of PVC substrate with no photoactive material added to it. Results are plotted over the results obtained from seedlings of the same species grown under the same conditions but using neutral filters to reduce the amount of light they grew under without changing the light spectrum (“Neutral Filters”), which are used a reference response under non-selective (neutral) filters.

**Figure 6.**
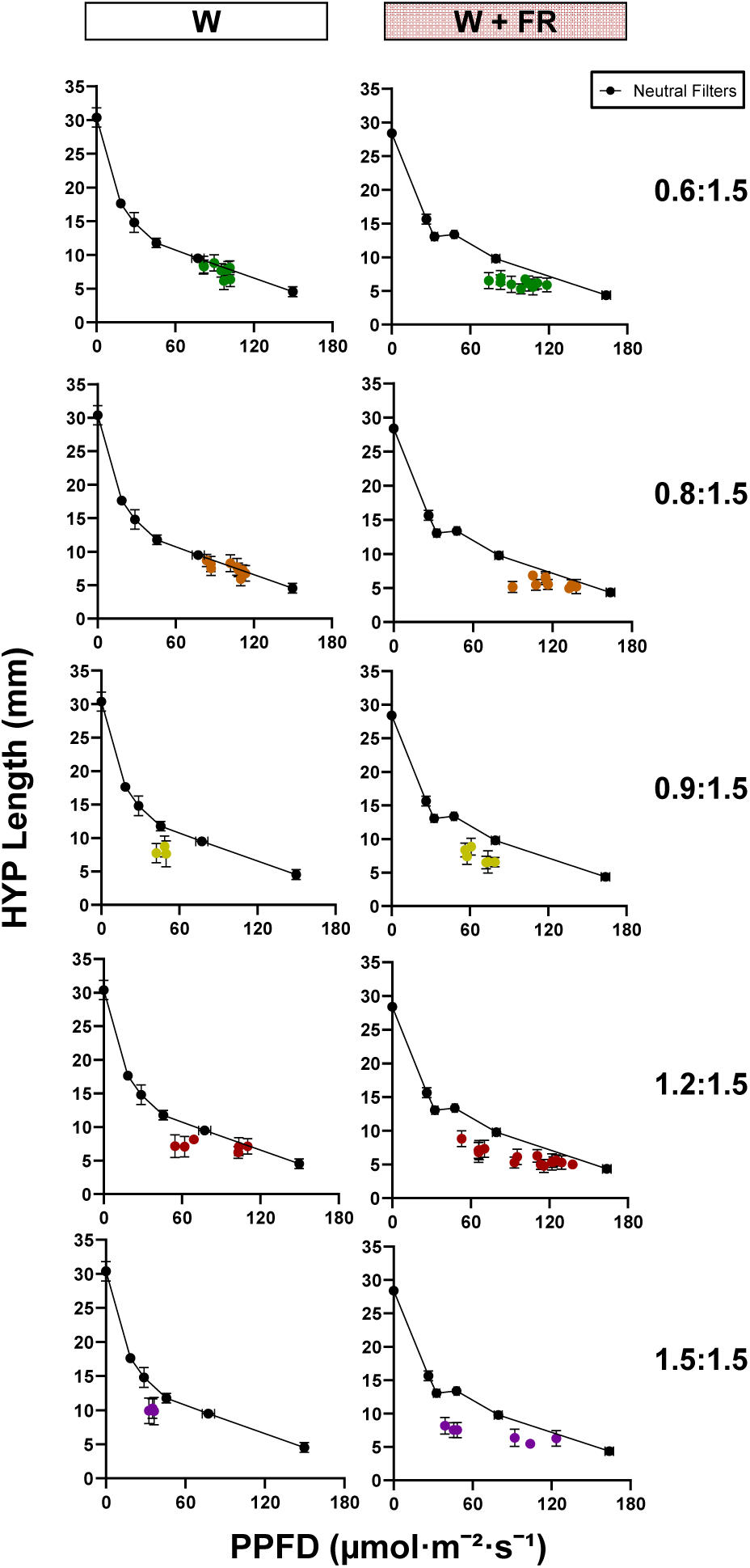
Hypocotyl elongation response of *C. hirsuta* under PTB7 – Th x IEICO 4F blended filters with different ratios of donor to acceptor material. Seedlings of *C. hirsuta* were grown for 7 days under filters made by blending PTB7 – Th (donor) x IEICO 4F (acceptor) materials in different ratios of donor to acceptor in W + FRc light treatment (R/FR ∼ 0.05). Graphs represent hypocotyl length relative to the Photosynthetic Photon Flux Density (PPFD). Values of hypocotyl length are the mean ±SD of three independent biological replicates. Each biological replicate is the mean of at least 15 seedlings. Values of PPFD correspond with three independent measurements taken for each replicate at the beginning of each experiment. Results are plotted over the results obtained from seedlings of the same species grown under the same conditions but using neutral filters to reduce the amount of light they grew under without changing the light spectrum (“Neutral Filters”), which are used a reference response under non-selective (neutral) filters.

## DISCUSSION

APV systems represent a promising strategy to reconcile the increasing demand for both food and energy production. However, their implementation inevitably modifies both light quantity and quality, two key environmental cues that strongly influence plant growth and development (Casal, 2012; Martinez-Garcia & Rodriguez-Concepcion, 2023). While most APV designs using opaque photovoltaic panels primarily reduce light intensity, semi-transparent organic photovoltaic (OPV) materials introduce an additional level of complexity by selectively filtering also specific regions of the light spectrum. Understanding how these spectral changes impact plant performance is therefore essential for optimizing APV systems.

Apart from both needing sunlight, plants and photovoltaic systems have distinct requirements in light quality and quantity. From these two aspects, generation of biomass in plants necessarily requires sufficient light energy to exceed the light compensation point. Therefore, one of the hypotheses when designing novel OPV materials is that they should minimally impact the photosynthetically relevant light to reach plants. However, light is a complex signal that also provides environmental information that can greatly impact on plant shape and form, architecture and/or developmental stages (e.g., transition from vegetative to reproductive stages). This informative aspect of the light is not trivial, as it might alter the amount and/or production of the organs that we normally harvest as food or feed. For instance, plants that lack all phytochromes, specialized in the R- and FR-sensing, exhibit impaired development, stalling at various stages despite being exposed to sufficient light for photosynthesis (Strasser et al., 2010). Despite the importance of light quality information for plant development, its impact on plant growth in opaque or semi-transparent OPV for APV systems has been poorly explored.

In this manuscript, we established a robust, sensitive and fast bioassay based on hypocotyl elongation to evaluate plant responses to OPV-filtered light. Among a few quick-and-easy-to-score morphological traits analyzed, hypocotyl length showed the highest dynamic range in response to light conditions in two model plant species, *A. thaliana* and *C. hirsuta*, confirming its suitability as a quantitative proxy for seedling growth under varying light environments (**Figure 1**). This is consistent with the well-established role of hypocotyl elongation as a highly plastic response integrating both light quantity and quality signals.

Our analyses using *A. thaliana* and *C. hirsuta* further validated the biological relevance of this bioassay. Interestingly, *C. hirsuta* exhibited reduced elongation (i.e., stronger light suppression of hypocotyl length) across light intensities (**Figures 2, S1**), supporting the idea that shade-tolerant species maintain growth under lower irradiance without triggering elongation responses (Morelli et al. 2021; Martinez-Garcia & Rodriguez-Concepcion, 2023). More importantly, these differences highlight the importance of considering a broad range of species displaying distinct light sensitivity when evaluating APV compatibility.

By comparing hypocotyl elongation of seedlings growing under the wavelength-selective absorption OPV filters and under neutral filters transmitting similar PPFD, we were able to disentangle effects of light quantity from those of light quality. This approach revealed that certain materials, such as PTB7-Th and D18, produced growth responses more similar to neutral shading (**Figures 3, 4**). Those OPVs that resulted in enhanced elongation indicated that the spectral composition of transmitted light rather than total light intensity alone in the 400-700 nm range, is also important to determine plant responses under OPV systems.

Further optimization of OPV performance requires balancing plant growth compatibility with photovoltaic efficiency. Combination of donor–acceptor blends indicated PTB7-Th and IEICO-4F as a suitable pair, and altering their ratio also modulates light transmission. Notably, increasing the donor proportion, which reduces overall light transmission, exhibited growth patterns similar to or even shorter than those under neutral filters (**Figures 5, 6**), suggesting the advantage of single or combinations of OPV materials that minimize distortion in the blue region.

The use of seedlings of small-size species in our assays offers clear and obvious advantages: it reduces the size of the filters, the required space and the time to conduct the experiments. Additionally, it allows to expand easily the number of OPVs that can be tested, lowering the cost of these analyses. More broadly, this work highlights the need to integrate plant photobiology into the design of next-generation APV systems. However, this approach assumes that seedling and adult plant respond similarly. Although in some cases this assumption holds true (e.g., hypocotyl length in seedlings vs. petiole length of adult leaves) future studies should extend these findings to adult plants of these same species but also to crop species. At these later stages, additional physiological traits, such as architecture and even yield, will be essential to fully evaluate the agronomic potential of OPV-based APV systems.

## EXPERIMENTAL PROCEDURES (still working on it)

### Plant material

For plant bioassays in this study, *A. thaliana* (Col-0 accession) and *C. hirsuta* (Oxford accession, Ox) were used (Molina-Contreras et al., 2019).

### Growth conditions for seedling bioassays

For hypocotyl elongation and cotyledon area and length assays, seeds were surface-sterilized and sown in Peri dishes with solid growth mdium without sucrose (0.5xMS-). Seeds were germinated below a window covered by the available OPV filters (6 x 6 cm) to ensure that only the transmitted light would impact plant material. This window allows to grow a maximum of 18 – 20 seeds of *A. thaliana* and *C. hirsuta* were sown in the middle of each Petri dish matching this window opening (maximum number of seeds that could be fitted in a 6 x 6 cm window). Sown seeds were put through stratification at 4°C in the dark for 3-4 days.

After stratification, the Petri dishes containing sown seeds were placed in the filter boxes and incubated for 7 days in dedicated plant growth chambers at 22°C and long day photoperiod conditions (16 h light, 8 h dark). White light (W) was emitted from LED light tubes providing a Photon Flux Density (PPFD) of a maximum of 150 µmolꞏm^-2^ꞏs^-1^ in the photosynthetically active radiation region (PAR, 400-700 nm) and a R:FR > 2.5. Shade conditions (W + FR) were simulated by supplementing W with FR provided by LEDs (www.philips.com/horti), resulting in a PPFD of a maximum of 170 µmolꞏm^-2^ꞏs^-1^ and a R:FR of 0.05.

### Light conditions for growth assays

Light fluence rates were measured using a Spectrosense2 meter associated with a 4-channel sensor (SkyeInstruments Ltd., www.skyeinstruments.com), which measures PAR (400-700 nm), and 10 nm windows of blue (467-473 nm), R (664-674 nm) and FR (725-735 nm) regions. Light spectra were generated using a Flame model Spectrometer with a Sony detector (FLAME-S; Ocean Optics) or a SpectraPen mini (PSI; https://psi.cz/).

Light measurements were taken by placing the sensor in place of each box in the plant growth chamber with the different filters covering the sensor.

### Measurements of seedling traits

Hypocotyl length was measured as indicated elsewhere (Pastor-Andreu et al., 2024). Briefly, seedlings were made to lay out flat on the plate media and digital images were taken and used to manually measure the length of each hypocotyl using the National Institutes of Health ImageJ software (http://rsb.info.nih.gov/).

For each experiment, 18 – 20 hypocotyl measurements were obtained for each tested filter and each species, which were then averaged. Each experiment was repeated 3 times in order to obtain 3 mean hypocotyl measurement values for each tested filter. Data was represented using GraphPad Prism (www.graphpad.com/).

## ACKNOWLEDGEMENTS

This work was funded by MICIN/AEI (MCIN/AEI/10.13039/501100011033) through the following grants: PLEC2022-009323 to MC-Q and JFM-G (Projects in strategic lines 2022, European Union NextGenerationEU/PRTR); PID2023-149395NB-I00 (ERDF/EU) to JFM-G; by the Generalitat Valenciana grant CIPROM/2024/41 to JFM-G.

## AUTHOR CONTRIBUTIONS

MC-Q and JFM-G conceived the original research plan and directed and coordinated the study. MV-T, MM-T and AAAT planned and performed the research and analyzed their respective data. JFM-G, MC-Q, MV-T and MM-T wrote the paper with revisions, contributions and/or comments from all authors.

## SUPPLEMENTARY FIGURES

**Figure S1.**
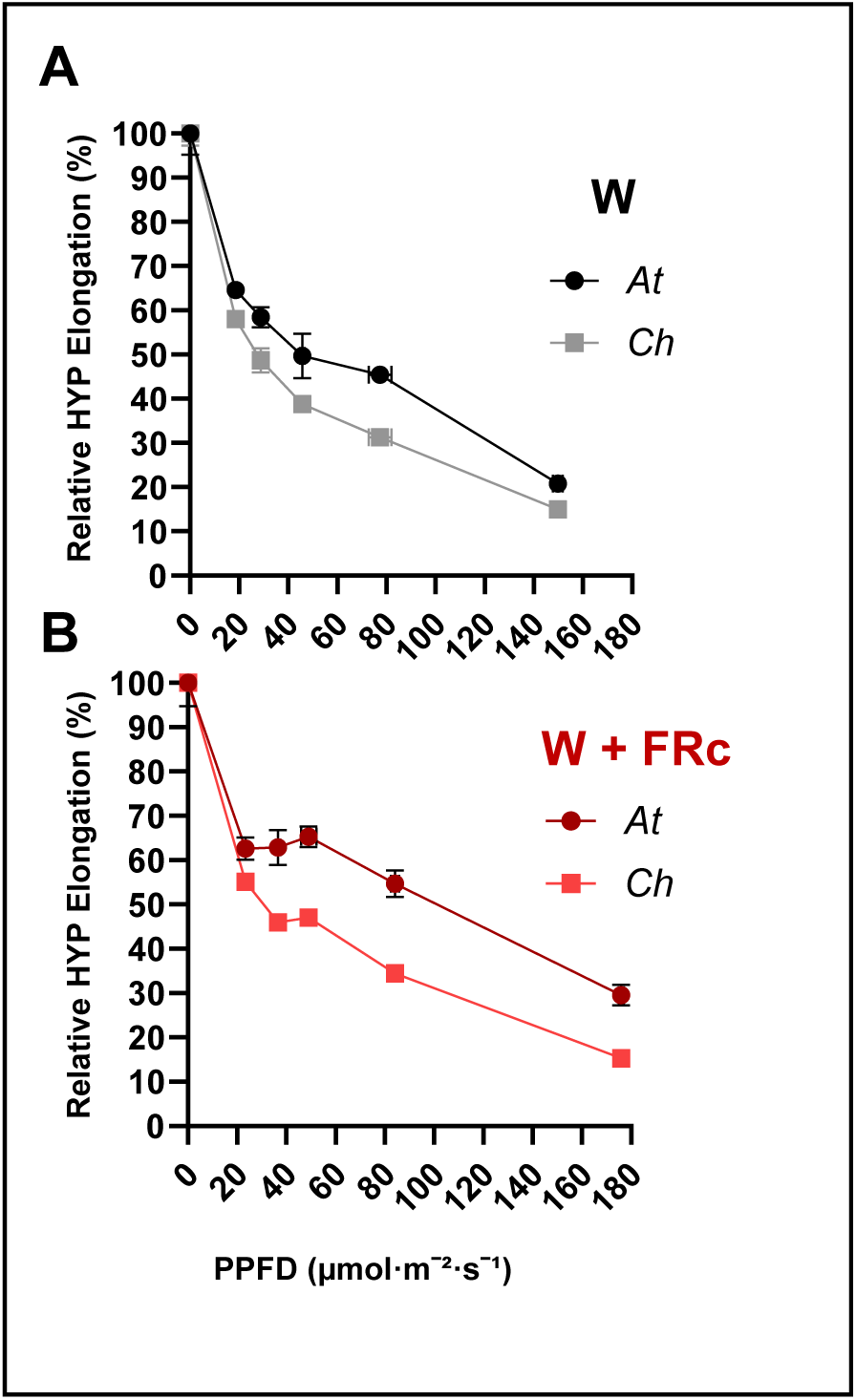
Relative hypocotyl elongation response of *A. thaliana* (*At*) and *C. hirsuta* (*Ch*) to different light quantity and quality conditions. These data were generated after processing those shown in Figure 2.

